# Polymorphisms in STING affect human innate immune responses to poxviruses

**DOI:** 10.1101/2020.05.28.121657

**Authors:** Richard B. Kennedy, Iana H. Haralambieva, Inna G. Ovsyannikova, Emily A. Voigt, Beth R. Larrabee, Daniel J. Schaid, Michael T. Zimmermann, Ann L. Oberg, Gregory A. Poland

## Abstract

We conducted a large genome-wide association study (GWAS) of the immune responses to primary smallpox vaccination in a combined cohort of > 1,600 subjects. We identified a cluster of SNPs on chromosome 5 (5q31.2) that were significantly associated (p-value: 1.3 × 10^−12^ – 1.5×10^−36^) with IFNα response to *in vitro* poxvirus stimulation. Examination of these SNPs led to the functional testing of rs1131769, a non-synonymous SNP in *TMEM173* causing an Arg-to-His change at position 232 in the STING protein—a major regulator of innate immune responses to viral infections. Our findings demonstrate important functional differences between the two alleles, where the major allele (R232) more effectively induces IFNα secretion. Molecular modeling of both alleles identified altered ligand binding characteristics between the two variants, providing a potential mechanism underlying differences in inter-individual responses to poxvirus vaccination. Our data demonstrate that possession of the H232 variant impairs STING-mediated innate immunity to poxviruses. These results clarify prior studies evaluating functional effects of genetic variants in *TMEM173* and provide novel data regarding genetic control of poxvirus immunity.

**Contribution to the Field:** Here we report that a single nucleotide non-synonymous polymorphism in the *TMEM173* gene encodes for a STING variant conferring a reduced IFN stimulated response compared to wild type. Our results suggest that, upon binding of the STING H232 variant to its ligand, activation of downstream signaling proteins is impaired, resulting in decreased production of IFNα and a weaker interferon-stimulated gene response. Molecular modeling indicates that the diminished functional activity of this variant is likely due to an altered physical structure of the STING protein. STING controls the innate, type I IFN response to double-stranded DNA and cyclic dinucleotides. Individuals with the H232 variant of STING have a much weaker innate immune response to vaccinia virus. Our data help resolve ongoing controversies regarding the role of genetic variants in STING function. Because STING plays an important role in our immune response to DNA viruses and bacteria, our results can be used to predict who will and will not respond to vaccines and treatments, and to design more effective vaccine candidates. Given the role of the STING protein in innate responses to DNA viruses and bacterial pathogens, these data may also be useful in developing novel treatment options for multiple infectious diseases.

## Introduction

Until its eradication in 1980, smallpox (caused by the variola virus) was a deadly, debilitating disease estimated to have killed hundreds of millions of individuals over the last two centuries alone (Fenner et al., 1988). Eradication was made possible by using vaccines based on vaccinia virus (Fenner et al., 1988). These live-virus vaccines elicited robust, long-lasting immunity in nearly all vaccine recipients (Crotty et al., 2003; Combadiere et al., 2004). Routine smallpox vaccination was halted in the United States before global eradication due to rare but serious adverse events, including death; however, poxviruses remain a public health issue for several reasons, including zoonotic poxvirus outbreaks (Di Giulio and Eckburg, 2004; Nagasse-Sugahara et al., 2004; Essbauer et al., 2010; Singh et al., 2012) and concerns regarding the release of variola virus as a biological weapon and novel poxviruses (Breman et al., 2003). The increasing use of poxviruses as platform vectors for other vaccines and therapeutics has also enhanced our need for a better understanding of poxvirus immunity. While highly effective, the smallpox vaccine has numerous contraindications as well as rare but serious, potentially life-threatening adverse reactions that limit its widespread use, if needed, in the population. Understanding how poxvirus immunity is controlled may assist in the development of safer yet still effective poxvirus-based vaccines and can provide insights into immunity to other DNA viruses/vaccines.

Although seroconversion rates after smallpox vaccination are high (>97%), antibody titers and cellular immune responses vary widely among recipients (Fenner et al., 1988; Kennedy et al., 2009a; Kennedy et al., 2009b; Kennedy et al., 2012a; Ovsyannikova et al., 2012). We have previously reported on a small subset of individuals who develop the classical vaccine take (i.e., response) but fail to mount vigorous adaptive immune responses (Kennedy et al., 2016). Previous research by our lab and others demonstrates that genetic polymorphisms are correlated with immune outcomes to multiple viral vaccines, including the smallpox vaccine (Crowe, 2007; Kennedy et al., 2012a; Kennedy et al., 2012b; Ovsyannikova et al., 2012). To move beyond statistical genetic associations, functional studies are also needed to elucidate the biologic mechanisms underlying these associations and link them to gain a better understanding of how genomic factors contribute to inter-individual variation in immune response.

We have previously reported the findings from the first genome-wide association study (GWAS) examining the association of single nucleotide polymorphisms (SNPs) with immune responses in a cohort of primary smallpox vaccines (Kennedy et al., 2012a; Kennedy et al., 2012b; Ovsyannikova et al., 2012). Here, we report a cluster of SNPs on chromosome 5 (5q31.2) that were significantly associated with IFNα response following *in vitro* stimulation of PBMCs with vaccinia virus. We report the results from the functional testing of rs1131769, which is a non-synonymous SNP in *TMEM173* that introduces an amino acid change from the arginine at position 232 (R232) to histidine (H232) in the STING protein—a major regulator of innate immune responses to viral and bacterial infections (Unterholzner et al., 2011; Cao et al., 2012; Ferguson et al., 2012; Georgana et al., 2018). Our results indicate that the H232 variant of STING is associated with a significant reduction in the IFNa response and that this effect is independent of the effect previously described for SNPs in the STING *HAQ* haplotype (Yi et al., 2013). We also report the results of molecular modeling and molecular dynamics (MD) simulations investigating differences in how the H232 and R232 variants interact with the signaling ligand. Overall, our study provides novel and important data regarding genetic control of poxvirus immunity in humans by linking specific genetic polymorphisms in *TMEM173* to differential STING pathway activation during innate immune responses (IFNα) to vaccinia virus. These results may also explain inter-individual variations in the innate immune response to other DNA viruses (e.g., HPV, VZV, HSV-1), which also stimulate the STING pathway, as well as the large number of bacterial pathogens that also activate STING.

## Materials and Methods

### Study cohorts

Two previously described study cohorts were combined for our analyses (Kennedy et al., 2012a; Kennedy et al., 2012b; Ovsyannikova et al., 2012). Briefly, the SD cohort consists of 1,076 Dryvax® recipients recruited in 2003–2006. The U.S. cohort consists of 1,058 ACAM2000® or Dryvax® recipients recruited in 2010–2013. For both cohorts, subjects had received their first (and only) smallpox vaccination between 1 and 4 years prior to study enrollment. All participants gave written informed consent for this study. Approval for all study procedures was obtained from the Institutional Review Boards of the Mayo Clinic (Rochester, MN) and the Naval Health Research Center (NHRC; San Diego, CA).

### Measurement of vaccinia-specific IFNα responses

Subject PBMC samples were cultured in the presence/absence of inactivated vaccinia virus (NYCBOH) at an MOI of 0.05. Vaccinia virus was inactivated using psoralen (5ug/ml) and long-wave UV light (Tsung et al., 1996). The full panel of cytokines included: IL-1β, IL-2, IL-4, IL-6, IL-10, IL-12p40, IL-12 p70, IL-18, IFNa, IFNb, and TNF-α. Interferon alpha (IFNα) production by vaccinia virus-stimulated PBMC samples was measured by commercial ELISA assay (PBL Biomedical Laboratories, Piscataway, NJ), as previously described (Kennedy et al., 2012b; Ovsyannikova et al., 2014a).

### Genotyping and fine mapping

DNA from all subjects was extracted using Gentra Puregene kits (Kennedy et al., 2012a). Genome-wide genotyping for the study cohort was performed as previously described (Kennedy et al., 2012a; Kennedy et al., 2012b; Ovsyannikova et al., 2012; Haralambieva et al., 2013). For the SD cohort (recruited in 2003–2006), subjects were genotyped with either the Illumina 550 array or the Illumina 650 array. Genotype quality control (QC) prior to imputation was conducted separately for each platform. QC for the Illumina 550 and 650 arrays involved removing monomorphic SNPs and those on the Y chromosome. We also removed all SNPs with a call rate <95%, and all subjects with a call rate <95%. SNPs were also excluded if they failed Hardy-Weinberg Equilibration (HWE) test p-value > 10^−5^. Genetic sex was verified by PLINK. Subjects in the U.S. cohort were recruited in 2010–2013 and genotyped with the Illumina Omni 2.5 array. For the Omni 2.5 array, mitochondrial SNPs, those on the Y chromosome, and monomorphic SNPs were removed. SNPs were required to have a call rate at least 99%, and subjects had a minimum call rate of 95%. No inconsistencies were found between reported sex and genetic-determined sex. Across these cohorts, a total of 2,062 subjects passed QC for genotyping.

The 1000 Genomes cosmopolitan samples (Build 37: African, AFR; American, AMR; Asian, ASN; European, EUR) served as a reference for SNP imputation. Observed SNPs were eliminated prior to imputation if they could not be converted to the forward strand or if more than one SNP mapped to a given position. The reference genome was filtered to exclude SNPs whose minor allele frequency (MAF) was < 0.005. The data were then phased using SHAPEIT (Delaneau et al., 2013) and imputed via IMPUTE2 (Howie et al., 2009). SNPs were included in analyses if their imputation dosage allele R^2^ was at least 0.3 and their MAF was at least 0.01. These GWAS QC restrictions resulted in a dataset with 6,210,296 SNPs for the HumHap550 array; 6,244,529 SNPs for the HumHap650 array; and 6,243,494 SNPs for the Omni 2.5 array.

Fine mapping on the chromosome 5 region was performed using a custom Illumina iSelect panel that included known SNPs in the following gene regions: *TMEM173, KCNN2, DNAJC18*, and *TRIM36* (the coding region, the intronic regions, and 10kb upstream and downstream in order to capture regulatory regions). We then identified all SNPs highly correlated (r2 > 0.9) with each of the target SNPs of interest based on the GWAS results. SNPs were excluded from the fine-mapping effort for the following reasons: low rank on the Illumina design score metric (indicating a low likelihood of successful genotyping); any Illumina error codes; previously genotyped SNPs; and monomorphic SNPs (based on HapMap and 1000 Genomes data). The resulting list of 2,406 SNPs were included on the Illumina iSelect panel.

The genotyping was performed in Mayo Clinic’s Clinical Genome Facility on 2,208 subjects: 2,011 subjects from the SD and U.S. cohorts; and 197 subjects used for quality control (55 negative controls, 48 trios of father/mother/child). 1,996 of these subjects passed all QC metrics filters (e.g., call rate at least 99%, duplicates removed, etc.). Of the genotyped SNPs, a total of 580 SNPs were used in the analysis (156 SNPs failed genotyping; 10 SNPs had call rates < 95%; 32 SNPs had HWE p-values < 10E^−6^, and 1,500 were monomorphic). For the Caucasian subgroup, an additional 126 SNPs were removed because they were monomorphic in that subgroup.

### Genetic ancestry and population stratification

Genotypes from the GWAS arrays were used to assign ancestry groups (i.e., Caucasian, African American, or Asian) to participants using the STRUCTURE software (Pritchard et al., 2000) and the 1000 Genomes reference data. Genetic ancestry proportions were estimated within cohorts and arrays (San Diego/550, San Diego/650, US/Omni 2.5), as previously described (Kennedy et al., 2012a; Kennedy et al., 2012b). An LD pruning process (Ovsyannikova et al., 2017) was utilized to ensure that the SNPs used for STRUCTURE and for sample eigenvectors were not drawn from small clusters within specific locations (Novembre and Stephens, 2008). Resulting SNPs were entered into to the STRUCTURE program (Pritchard et al., 2000), and participant ancestry was classified based on the largest ancestry proportion estimated by STRUCTURE.

Within ancestry groups, eigenvectors were estimated for population-stratification purposes. SNPs with a MAF < 0.01 and those with a HWE p-value < 0.001 were excluded, as were insertion/deletions (INDELS). The remaining SNPs were pruned according to the following variance inflation factors: window size of 50 kilobases; step size of 5; and variance inflation factor threshold of 1.05. SmartPCA was used to create the eigenvectors (Price et al., 2006) following the procedures implemented in EIGENSTRAT software. Eigenvectors were included as potential covariates if they had a Tracy-Widom p-value < 0.05.

### Selection of covariates to adjust for potential confounders

For analysis purposes, the immune-response trait of interest (IFNα secretion) was calculated by first computing the difference of the mean stimulated and unstimulated values and then transforming to a normal distribution using normal quantiles. In order to combine data from the two cohorts, potential confounder effects for each ancestry group and cohort were adjusted by linear regression models as described (Ovsyannikova et al., 2017). Categorical variables with a very large number of categories were binned using hierarchical clustering. This was achieved by using hierarchical clustering on the estimated regression coefficients for the different categories while binning categories with similar regression coefficients. Categorical variables were included in regression models by using indicator variables for categories, treating the most common category as baseline. Residuals from the linear models were used as the primary adjusted traits for GWAS analyses.

### GWAS analysis strategy

In order to maximize the power to detect SNPs associated with smallpox vaccine immune response phenotypes, data across genotyping arrays and the two cohorts was pooled after preparing the data as described above. The pooled analyses were then performed using the adjusted traits described above in a regression analysis, along with an indicator of cohort as an additional adjusting covariate. Because the largest ancestry group was Caucasian, we restricted our pooled analysis to the Caucasian subjects (n=1,605). Multiple testing was controlled for by using the standard p-value threshold (p-value< 5×10^−8^) to determine genome-wide statistical significance (Pe’er et al., 2008; Manolio, 2010). Statistical analyses were performed with the R statistical software and PLINK (Purcell et al., 2007).

### Generation of stable BJAB cell lines

BJAB stable cell lines, each expressing one of the rs1131769 variants of interest, were created using custom suCMV promoter-based lentivectors containing a Blasticidin resistance gene (GenTarget Inc.; San Diego, CA). Transduction with lentiviral particles was performed at MOI of 10 in the presence of Polybrene (8 μg/mL), and stable cell clones were selected for using Blasticidin (InvivoGen; San Diego, CA) at 10 μg/mL.

### Protein expression and phosphorylation by western blotting

Protein expression and phosphorylation (for IRF3 and TBK1) was assessed in transiently transfected HEK 293 T cells and lentivirus-transduced stable BJAB cell lines, expressing the STING alleles of interest. For HEK 293 T cells, transfection was performed with 20 ng of the STING plasmid constructs (InvivoGen; San Diego, CA, or DNA2.0, Inc.; Newark, CA) and with 20 ng cGAS, using Lipofectamine LTX and PLUS™ Reagent (Invitrogen; Carlsbad, CA), according to the manufacturer’s instructions. For all experiments, cells were incubated overnight in antibiotic-free medium and then stimulated with 2’3’cGAMP (20 μg/mL for the HEK 293 T cells and 100 μg/mL for the BJAB cells) for different time periods (15 min, 30 min, 1 hr, 2hrs, and 4hrs). The cells were lysed in RIPA lysis buffer (Sigma) containing protease and phosphatase inhibitors. Lysates were centrifuged at 16,000 RPM and 4°C for 20 mins. Protein concentrations were quantified using the Pierce BCA Protein Assay Kit (ThermoFisher Scientific; Minneapolis, MN), and equal protein amounts (2 to 3μg) were used for western blot analysis. Laemmli buffer (Bio-Rad; Hercules, CA) with β-mercaptoethanol was added to the samples, and the lysates were denatured by incubating at 95°C for 5 min and were centrifuged at 16,000 RPM for 1 minute. Samples were loaded onto 4–20% Criterion™ gels (Bio-Rad; Hercules, CA), and then proteins were transferred to Trans-Blot® Turbo Midi PVDF membranes (Bio-Rad; Hercules, CA) using the Trans-Blot® Turbo™ Transfer System (Bio-Rad; Hercules, CA). Blots were blocked with 3% BSA and probed overnight (at 4°C) with primary rabbit anti-STING, anti-TANK-binding kinase 1 (TBK1), anti-pTBK1, anti-interferon regulatory factor 3 (IRF3), and anti-pIRF3 antibody (all from Cell Signaling Technologies; Beverly, MA), or mouse anti-alpha tubulin antibody (Abcam; Cambridge, MA) for loading control. Membranes were washed and incubated for 1 hr at room temperature with the appropriate HRP-labeled pre-absorbed goat anti-rabbit or anti-mouse secondary antibodies (Santa Cruz Biotechnology, Inc.; Dallas, TX). The membranes were washed, developed using Clarity Western ECL Substrate Solution (Bio-Rad; Hercules, CA) for 10 min, and imaged using the ChemiDoc™ Touch Gel Imaging System (Bio-Rad; Hercules, CA).

### qPCR

STING allele-expressing BJAB cells were plated at 300,000 cells/mL, 0.5 mL/well in a 24-well plate. 50 ug/mL of 2’3’ cGAMP or ddH2O were added to stimulated and mock-stimulated wells, respectively. Stimulated and control cells were harvested at the indicated times after 2’3’ cGAMP stimulation and centrifuged 5 min at 5,000 rpm in microcentrifuge tubes. Supernatants were collected and frozen for ELISA analysis. Cells were resuspended in 200 uL RNAProtect (Qiagen; Valencia, CA) and frozen at −20°C until RNA extraction and qPCR. Total RNA was extracted from cells using Qiagen RNeasy Plus mini kits according to the manufacturer’s instructions, and RNA concentrations were normalized between samples.

Random-primer reverse transcription was done using RT2 First Strand kits (Qiagen; Valencia, CA), including a genomic DNA removal treatment, according to the manufacturer’s instructions. qPCR was then done on each sample using the Qiagen RT2 SYBR Green/ROX qPCR Mastermix system using the following primers (Voigt and Yin, 2015): IFN-α, 5’-AAATACAGCCCTTGTGCCTGG-3’and 5’-GGTGAGCTGGCATACGAATCA-3’; IFN-β, 5’-AAGGCCAAGGAGTACAGTC-3’ and 5’-ATCTTCAGTTTCGGAGGTAA-3’; IFN-λ1, 5’-CGCCTTGGAAGAGTCACTCA-3’; IFN-λ1 5’-GAAGCCTCAGGTCCCAATTC-3’; b-actin, 5’-AAAGACCTGTACGCCAACAC-3’; b-actin 5’-GTCATACTCCTGCTTGCTGAT-3’; STING, Commercial Qiagen RT2 qPCR Primer Assay for Human TMEM173, MxA, Commercial InvivoGen IFNr qRT-Primer set, hOAS1-F and hOAS1-R; OAS1, Commercial InvivoGen IFNr qRT-Primer set, hMX1-F and hMX1-R. Quantitative PCR was done using an ABI ViiA-7 machine at the standard qPCR conditions starting with incubation at 95°C for 10 min, followed by 40 cycles of 95°C for 15 seconds and 60°C for 1 minute. Ct values were normalized to β-actin levels and unstimulated controls by the standard 2ΔΔCT method.

### Promoter reporter assays

The promoter reporter assays were performed in HEK 293 T cells, stably expressing one of the STING alleles of interest (for rs1131769 – WT/R232 and H232) under blasticidin selection (Antoniak et al., 2013). We used pNiFty2-IFNB-SEAP and pNiFty2-56K-SEAP promoter-reporter plasmids (InvivoGen; San Diego, CA), encoding the INFβ minimal promoter and the ISG-56K promoter, respectively. Co-expression with constitutively activated IRF3 (or IRF7) leads to promoter induction measured by the inducible expression of the secreted embryonic alkaline phosphatase (SEAP) reporter gene. Promoter assays were performed as previously described (Antoniak et al., 2013) but with some modifications. Briefly, 2.5 × 10^4^ cells per well (stably expressing STING alleles of interest under blasticidin selection) were cultured overnight in 96-well plates in antibiotic-free medium (DMEM [Gibco Invitrogen Corporation; Carlsbad, CA]), containing 10% fetal bovine serum (FBS, HyClone; Logan, UT). On the following day, cells were transfected with Lipofectamine® LTX (Invitrogen; Carlsbad, CA), according to the manufacturer’s protocol, using a constant amount of reporter plasmid (100 ng of either pNiFty2-IFNB-SEAP or pNiFty2-56K-SEAP per well), 0.2 μL PLUS™ Reagent (Invitrogen; Carlsbad, CA) per well, and 0.25 μL Lipofectamine LTX per well. After overnight incubation, the medium was switched to Opti-MEM (Gibco Invitrogen Corporation; Carlsbad, CA), and cells were stimulated with one of two STING ligands: 2’3’ cGAMP (100 μg/ml), or inactivated vaccinia virus (MOI of 10) at 37°C for different time periods. Promoter induction was measured by the SEAP reporter secretion (quantified at 620nm following addition of Quanti-Blue™ media, Invivogen, per the manufacturer’s instructions).

### ELISA measurement of secreted type I and type III IFNs

IFNα and IFNλ production by 2’3’ cGAMP-treated STING-transduced BJAB cells were measured in triplicate using commercial sandwich ELISA assay sets (IFNα: VeriKine-HS™ Human Interferon Alpha All Subtype ELISA Kit, PBL Assay Science; Piscataway, NJ, and IFNλ: Human IL-29/IL-28B [IFN-lambda 1/3] DuoSet ELISA set, R&D Systems; Minneapolis, MN) according to the manufacturer’s instructions. Standard protein samples were diluted in cell culture media for accurate standard curve construction and calculations. Recombinant IFNs were used as positive controls while cell culture media served as the negative control. Biological duplicate samples from each timepoint were assayed in duplicate.

### Molecular modeling

The atomic structure of the cyclic dinucleotide binding domain of STING has been experimentally solved (Huang et al., 2012). As is common for crystallographic structures, mobile loops were not resolved in these structures. To initially place residues within these mobile loops, we used the SwissModel server (Biasini et al., 2014) and template PDB structures 4QXP (Gao et al., 2014) (open conformation with inhibitor bound) and 4F5Y (Shang et al., 2012) (closed conformation with cdGMP bound). Mutations present in each template were reverted to WT amino acids according to the UniProt sequence of the canonical transcript (Q86WV6-1). Simulations were run for the apo (un-liganded), cdGMP, and cGAMP ligand states.

We used NAMD (Phillips et al., 2005) and the CHARMM27 with the CMAP (Mackerell et al., 2004) force field for Generalized Born implicit solvent molecular dynamics (isMD) simulations using previously optimized conditions (Sethi et al., 2018) that included the following: 1) an interaction cutoff of 15Å; 2) strength tapering (switching) starting at 12Å; 3) a 1fs simulation time step with conformations recorded every 2ps; 4) an initial conformation that was energy minimized for 20,000 steps; and 5) heating to 300K over 300ps via a Langevin thermostat. From each of the 12 conditions (two initial conformations, two alleles, and three ligand states), 100ns of simulation trajectory was generated and the final 70ns analyzed. Three additional and independent 20ns replicates for each condition were generated using the same procedure. All trajectories were aligned to the initial R232 closed conformation using C^α^ atoms. Trajectories were then evaluated using multiple metrics, including C-alpha Root Mean Squared Deviation (RMSD), Root Mean Squared Fluctuation (RMSF), Principal Component (PC) analysis, alignment-free distance difference matrix (Holm and Sander, 1993; Grant et al., 2006; Rashin et al., 2014), and distance monitors across the ligand binding site. We quantified variance of atomic C^α^ - C^α^ (Figure 7A) distances using median absolute difference (MAD). Analysis was performed using custom scripts on the Bio3D R package (Grant et al., 2006) and VMD (Humphrey et al., 1996).

## Results

### Overview of Cohorts and IFNα Response

We conducted a GWAS in the initial cohort of 1,076 primary smallpox vaccine recipients to identify regions of interest for fine mapping (Kennedy et al., 2012a; Kennedy et al., 2012b; Ovsyannikova et al., 2012). We then recruited a second cohort of primary smallpox vaccine recipients (n=1,058) and imputed additional SNPs, as described in Methods. The cohorts were combined, and a final study sample of 1,605 Caucasian subjects was used in the analysis (See Supplemental Table 1 for demographic information). Our original intent was to determine whether or not there were genetic polymorphisms associated with markers of vaccine-induced cellular immunity. We previously reported on the cytokine response following *in vitro* stimulation of these vaccinated subjects’ PBMCs with inactivated vaccinia virus (Umlauf et al., 2011; Haralambieva et al., 2013; Ovsyannikova et al., 2014b). VACV was inactivated in order to minimize the immunomodulatory effect of poxvirus-encoded proteins and to allow full development of the cytokine response. Included in our panel were multiple cytokines (e.g., IL-1b, IL-6, IFNα) produced during innate antiviral responses, as well as cytokines associated with adaptive immune function. Of note, in this GWAS analysis, the only signal reaching genome-wide significance was associated with IFNα secretion, which is reflective of an innate response to poxvirus infection rather than a T cell response due to vaccine-induced immunity. This result suggests that our findings are broadly applicable to poxvirus immunity and not just smallpox vaccination.

The GWAS analysis identified an exceptionally strong signal on chromosome 5 (Figure 1A and 1B). The locus-zoom plot in Figure 1C depicts the SNPs with the strongest statistical association with IFNα response. We used a genome-wide threshold of p-value <5 × 10^−8^ to establish statistical significance. Further details on the SNPs meeting this threshold are provided in Table 1. Our combined study cohort had a median IFNα secretion level of 126 pg/mL (IQR: 48.6-229.6) in PBMC cultures after vaccinia virus stimulation. As illustrated in Figure 2, TT homozygotes (H232 STING) had a median IFNα response of 17.7 pg/mL, while individuals homozygous for the R232 STING allele (CC genotype) had an 8-fold higher response (143.6 pg/mL). Heterozygotes had an intermediate phenotype.

**Figure 1.**
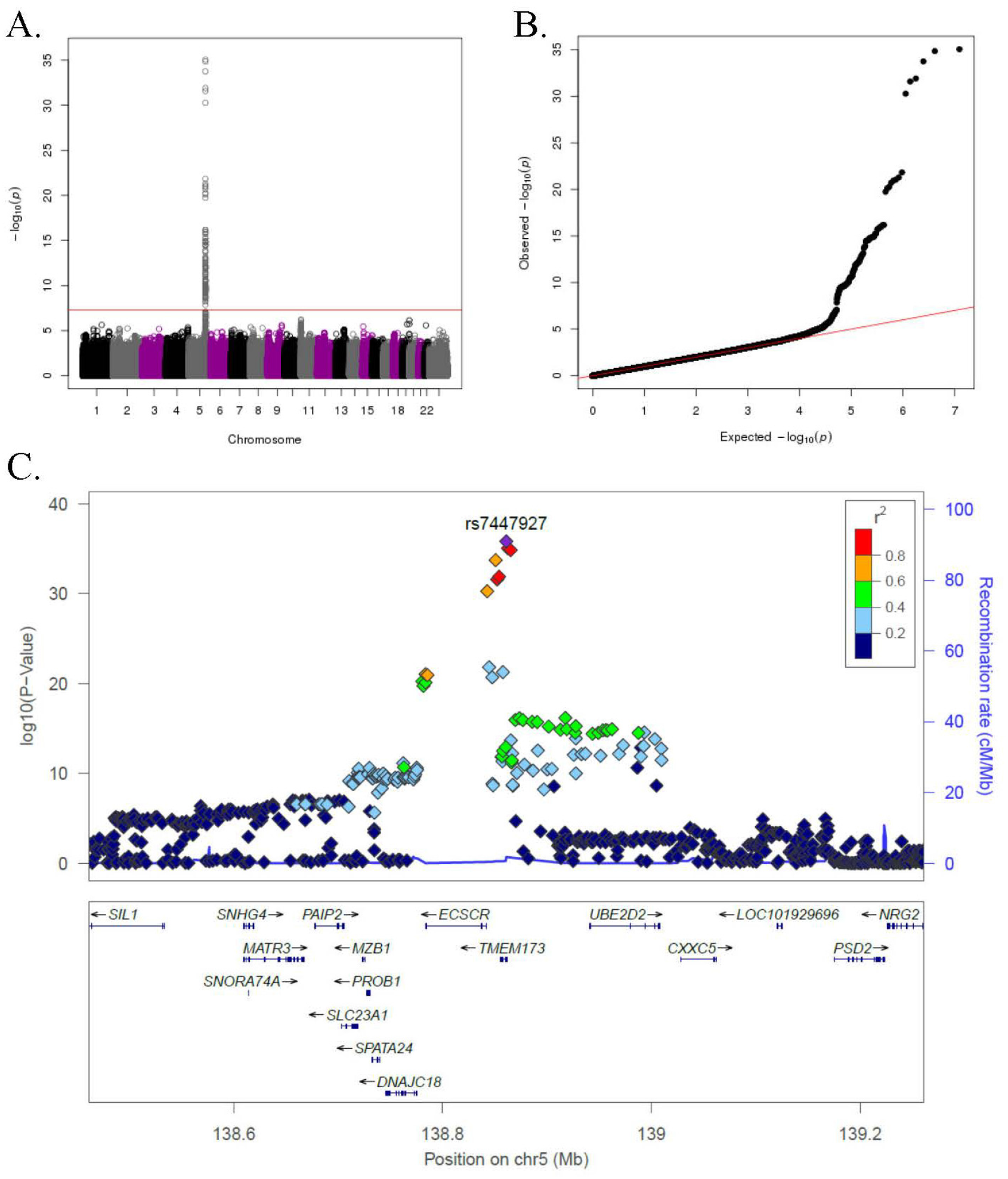
GWAS genotyping results in the combined smallpox vaccine recipient cohort. **A)** Manhattan plot indicating SNPs associated with IFNα response. **B)** QQ plot of genome-wide p-values. **C)** Locus-zoom plot depicting region on chromosome 5 with the strongest association signal. SNP LD is shown in color. The name and location of each gene is shown at the bottom of the panel.

**Figure 2.**
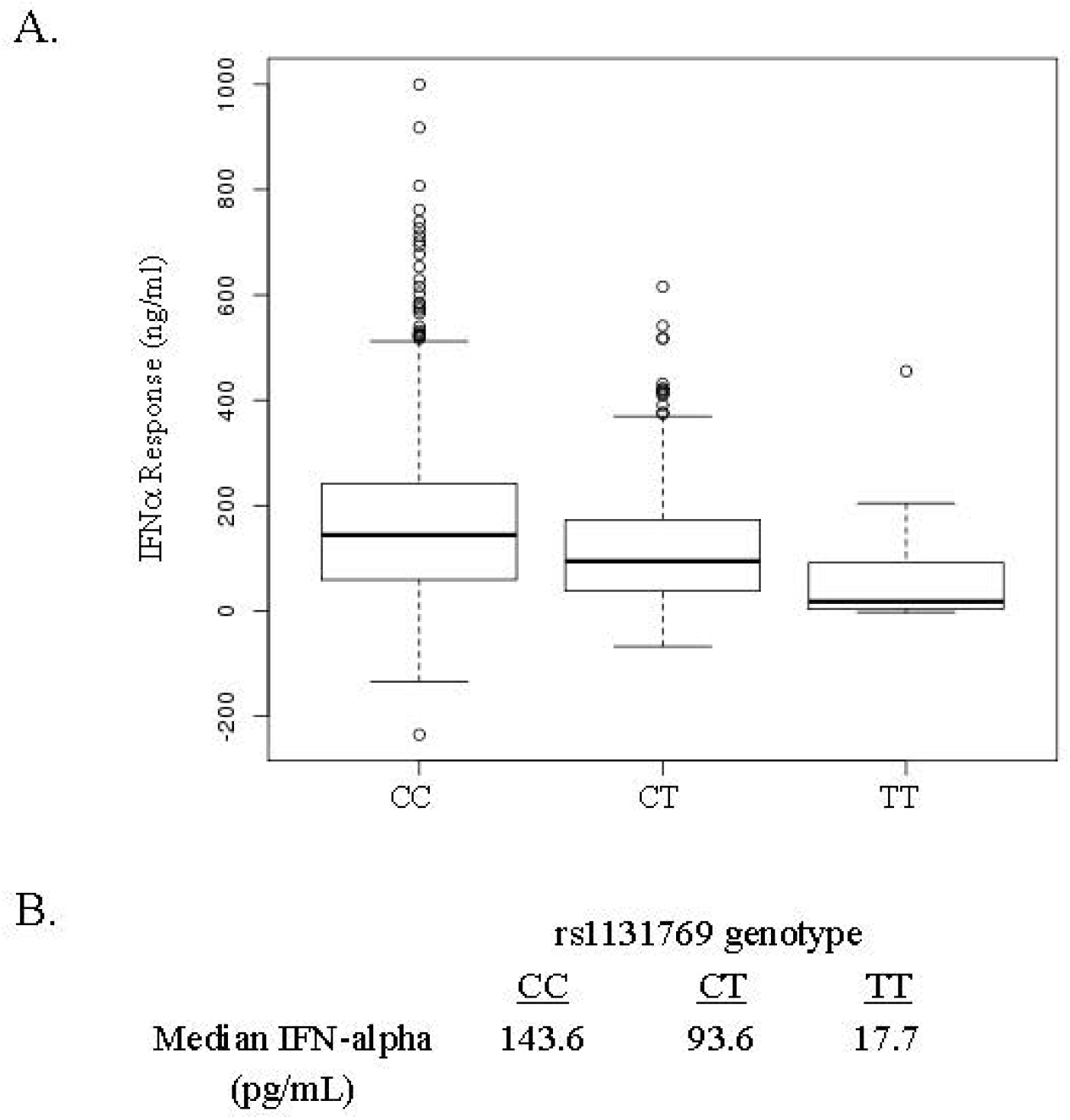
IFNa response in smallpox vaccine recipients displays a dose-dependent association with rsll31769. **A)** Box and whisker plots for major allele homozygotes (CC), heterozygotes (CT), and homozygous minor allele (TT) subjects. The C allele corresponds to the R232 STING variant and the T allele corresponds to the H232 STING variant. **B)** Median IFNa response (pg/mL) by rs 1131769 genotype group.

**Table 1.** Top SNPs significantly associated with vaccinia virus-specific IFNα secretion.

| SNP | Chromosome Location | Gene | SNP Function | Gene Location | p-value | Minor allele | Major allele | MAF |
| --- | --- | --- | --- | --- | --- | --- | --- | --- |
| rs7447927 | 138861146 | TMEM173 | protein-coding | synonymous | 1.49E-36 | C | G | 34.7% |
| rs13166214 | 138862744 | TMEM173 | protein-coding | 5'upstream | 8.92E-36 | A | G | 35.3% |
| rs7444313 | 138865423 | TMEM173 | protein-coding | 5'upstream | 1.41E-35 | G | A | 34.5% |
| rs13181561 | 138850905 | TMEM173 | protein-coding | 3'downstream | 1.81E-34 | G | A | 30.0% |
| rs55792153 | 138854203 | TMEM173 | protein-coding | 3'downstream | 1.26E-32 | A | C | 34.2% |
| rs13153461 | 138852369 | TMEM173 | protein-coding | 3'downstream | 2.61E-32 | G | A | 31.4% |
| rs9716069 | 138842818 | ECSCR | protein-coding | 5'upstream | 5.32E-31 | T | A | 31.3% |
| rs28419191 | 138844599 | ECSCR | protein-coding | 5'upstream | 1.50E-22 | T | C | 13.2% |
| ***rs1131769*** | ***138857919*** | ***TMEM173*** | ***protein-coding*** | ***missense*** | ***5.25E-22*** | ***T*** | ***C*** | ***14.0%*** |
| rs11954057 | 138783832 | RNU5B-4P | pseudo | 3'downstream | 8.99E-22 | C | G | 32.5% |
| rs36137978 | 138785565 | ECSCR | protein-coding | 5'upstream | 1.13E-21 | C | A | 31.6% |
| rs10875554 | 138847652 | ECSCR | protein-coding | 5'upstream | 1.99E-21 | A | C | 15.4% |
| rs6596479 | 138780599 | RNU5B-4P | pseudo | 5'upstream | 5.63E-21 | C | T | 31.9% |
| rs7446197 | 138783734 | RNU5B-4P | pseudo | 3'downstream | 7.51E-21 | A | G | 33.8% |
| rs10463977 | 138781765 | RNU5B-4P | pseudo | 5'upstream | 1.80E-20 | C | T | 32.4% |
| rs2434576 | 138917674 | UBE2D2 | protein-coding | 5'upstream | 6.57E-17 | G | A | 30.8% |
| rs34530489 | 138873627 | LOC642262 | pseudo | gene | 7.00E-17 | G | A | 31.4% |
| rs35779874 | 138869847 | LOC642262 | pseudo | 5'upstream | 1.11E-16 | A | G | 31.2% |
| rs7378724 | 138876953 | LOC642262 | pseudo | gene | 1.13E-16 | G | A | 30.9% |
| ***rs78233829*** | ***138857925*** | ***TMEM173*** | ***protein-coding*** | ***missense*** | ***3.16E-13*** | ***G*** | ***C*** | ***17.6%*** |
| ***rs11554776*** | ***138861078*** | ***TMEM173*** | ***protein-coding*** | ***missense*** | ***1.05E-12*** | ***T*** | ***C*** | ***16.5%*** |
| ***rs7380824*** | ***138856982*** | ***TMEM173*** | ***protein-coding*** | ***missense*** | ***1.25E-12*** | ***T*** | ***C*** | ***17.7%*** |

MAF: Minor allele frequency. Bold, italics – SNP studied in this report. Bold – SNPs in HAQ STING haplotype.

Several of the SNPs significantly associated with IFNα secretion were located in *TMEM173,* which encodes for STING—a pattern-recognition receptor mediating type I IFN responses to cyclic dinucleotides and double-stranded DNA. STING has previously been shown to play an important role in the innate immune response to poxviruses (Unterholzner et al., 2011; Dai et al., 2014).

A number of additional non-synonymous polymorphisms potentially affecting STING function have previously been identified, including R71H (rs11554776), G230A (rs78233829), R293Q (rs7380824), and R232H (rs1131769) (Jin et al., 2011; Yi et al., 2013). In order to determine which SNPs in the haplotype contributed to the response, while accounting for correlation among SNPs, we used haplo.stats software in R to compute the haplotype frequencies for these four SNPs: rs11554776; rs78233829; rs1131769; and rs7380824 (Schaid et al., 2002).

The haplotype frequencies were very close between the U.S. and San Diego (SD) cohorts; therefore, we proceeded to use the combined sample to evaluate the association of haplotypes with IFNα response. The results presented in Table 2 illustrate two key points. First, when comparing haplotype CCTC (R232H) versus the baseline (WT, wild type) haplotype CCCC, which focuses on the effect of the T allele for rs1131769 while controlling for the effects of the other three SNPs, there was a statistically significant (p ≪2E^−16^) decrease in IFNα response. Second, when comparing haplotype TGCT versus the baseline haplotype CCCC, which focused on rs7380824 (R293Q) while controlling for rs1131769, there was also a statistically significant (p ≪2E^−16^) decrease in IFNα response. The other two haplotypes (CCCT and CGCT) also had a decreased IFNα response, but the power to detect the effect of these haplotypes was limited due to their rarity. These analyses suggest that there are two independent effects on IFNα response—one from rs7380824 and another from rs1131769. To further explore this, we computed the dose of the minor allele for each of the four SNPs and performed backward regression. The two SNPs rs7380824 and rs1131769 remained in the model (each with p-value < 2E^−16^), illustrating that each SNP is strongly associated with IFNα response after adjusting for the other SNP. The other two SNPs, rs78233829 and rs11554776, were not statistically significant and were both strongly correlated with rs7380824 (Pearson correlations > 0.95).

**Table 2.** The Effect of rs1131769 on IFNα Response is Independent of the Effect Mediated by the HAQ Haplotype.

| Term in Model | Regression Coefficient** | Standard Error of Coefficient | p-value*** | Haplotype Frequency |
| --- | --- | --- | --- | --- |
| **Intercept** | 0.27 | 0.038 | 6.54E-13 |  |
| **Cohort US vs. SD** | 0.034 | 0.044 | 0.4360 |  |
| **Haplotypes\*** |  |  |  |  |
| CCCC | Baseline |  |  | 0.683 |
| CCTC (*H232*) | -0.542 | 0.045 | <2E-16 | 0.139 |
| CCCT (*Q293*) | -0.045 | 0.31 | 0.88 | 0.002 |
| CGCT (*AQ*) | -0.285 | 0.15 | 0.052 | 0.011 |
| **TGCT (*HAQ*)** | -0.407 | 0.042 | <2E-16 | 0.164 |

\*TMEM173 SNPs (haplotype) from left to right: rs11554776 (encodes amino acid/AA change at position 71) —rs78233829 (encodes AA change at position 230) — rs1131769 (encodes AA change at position 232) — rs7380824 (encodes AA change at position 293). In **bold** are designated the minor alleles defining the respective haplotypes, and in *italics* are designated the commonly used names of the haplotypes (that are based on encoded amino acid/acids and their position).
\*\* Regression coefficients from regression analysis of TMEM173 SNP haplotypes with IFNα response. Show the direction and magnitude of the estimated haplotypic effect on IFNα response compared to the haplotype (CCCC) with the greatest population frequency.
\*\*\*P-value from regression analysis.

The major allele of rs1131769 corresponds to a STING protein with Arg at position 232 (R232) and the minor allele encoding for His at position 232 (H232). The frequency of the minor T allele was 14.0% in our study cohort, which is close to the 12.7% reported in dbSNP. We constructed expression plasmids for these *TMEM173* variants and, using lentiviral constructs, created cell lines expressing each allele of rs1131769 in order to carry out experimental studies to elucidate functional differences between the rs1131769 STING variants.

### Promoter Activity of rs1131769 Variants

Two plasmids expressing the secreted embryonic alkaline phosphatase (SEAP) reporter gene, under control of either the INFβ promoter or the interferon stimulated gene (ISG)-56K promoter, were used to measure ligand-stimulated promoter activity in HEK293T cells expressing either R232 or H232 STING (Figure 3). Both 293T variant cell lines expressed high levels of STING mRNA (Figure 4C) with no significant difference between alleles. Upon 2’3’ cGAMP stimulation, IFNβ promoter induction was significantly higher at 10 hrs post-stimulation in R232 cells compared to cells expressing H232 (Figure 3A, p=0.02). Similarly, we observed statistically significant higher induction of the ISG-56K promoter activity in R232 upon stimulation with either 2’3’ cGAMP at 4 hrs and 8 hrs post-stimulation (Figure 3B, p=0.006 and p=0.004, respectively) or inactivated vaccinia virus at 8 hrs post-stimulation (Figure 3C, p=0.002).

**Figure 3.**
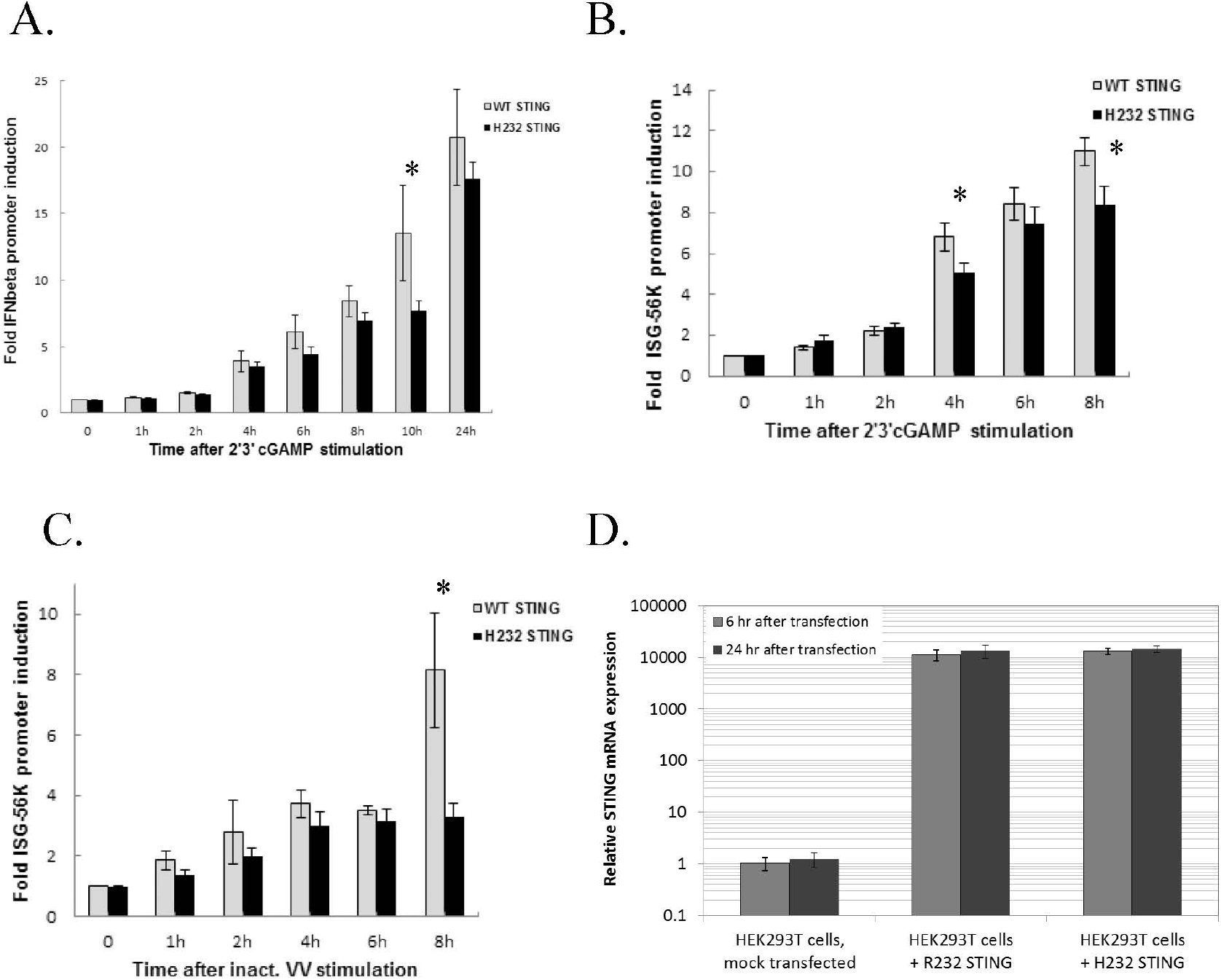
IFNβ and ISG-56K Promoter Activity is Greater in HEK 293T Cells Stably Expressing the R232 STING Variant. **A)** IFNβ promoter induction following cGAMP stimulation. **B)** ISG-56K promoter activity after cGAMP stimulation. **C)** ISG-56K promoter activity after stimulation with inactivated vaccinia virus (inact. VV). **D)** STING expression in HEK293T stably expressing R232 or H232 STING. * significant differences detected, see text for p-values.

**Figure 4.**
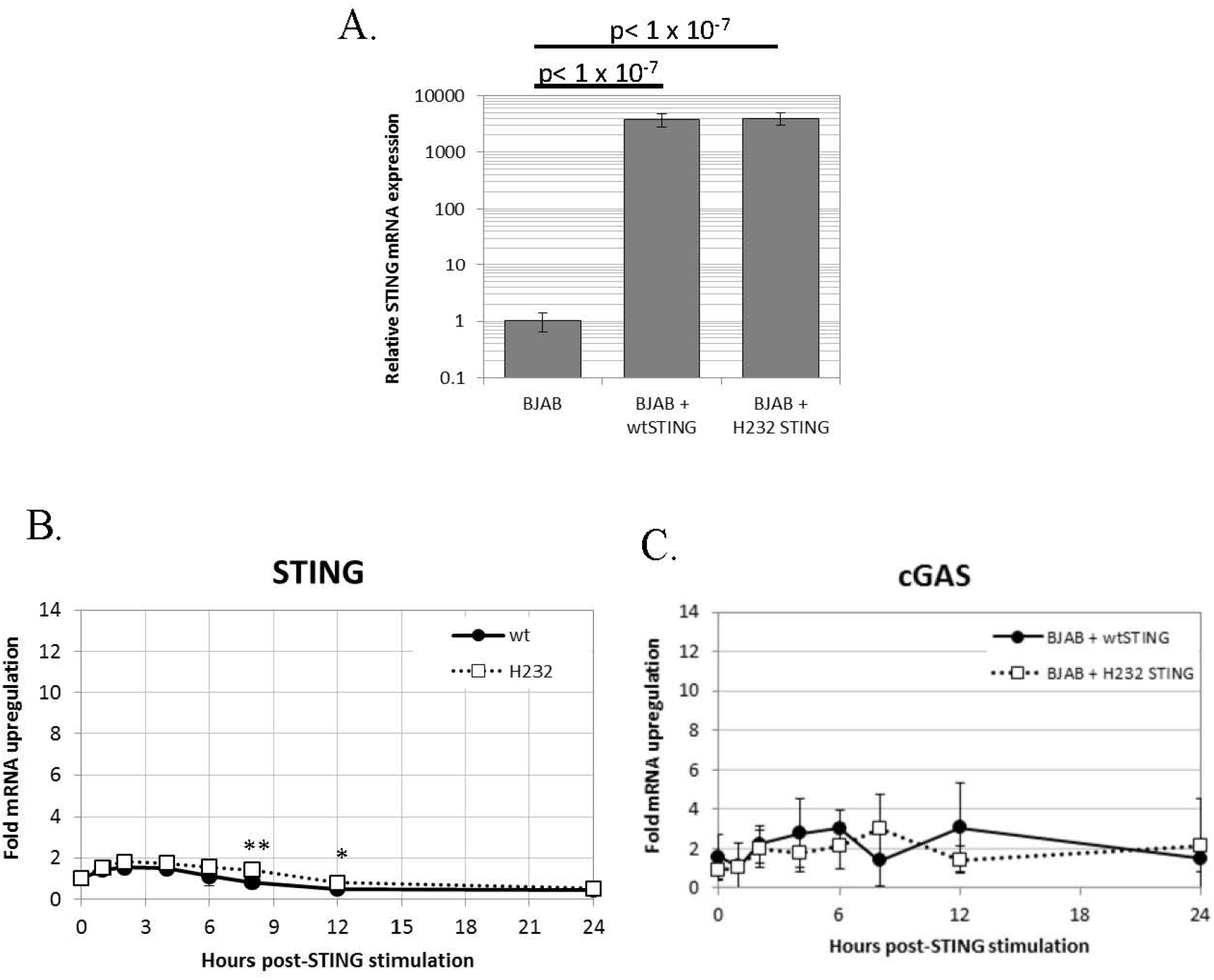
Stable transfection of H232 or R232 STING into BJAB cells results in high level expression that is stable upon cGAMP stimulation. **A)** Untransduced and stably transduced BJAB cell lines expressing STING alleles were harvested and assayed for STING mRNA using quantitative PCR. Values are shown as fold-levels relative to normal STING expression in untransduced BJAB cells. Data represent means and standard deviations of four biological replicates, assayed with technical duplicates. **B)** Time course of TMEM173 expression in 2’ 3’ cGAMP-stimulated (50ug/ml) in transduced BJAB cells stably expressing STING variants. **C)** Time course of cGAS expression in stable BJAB transfectants stimulated with cGAMP. Values are presented as fold-increases over mock-treated cells, normalized to β-actin loading controls. Data points are the average of 8 replicates. * p< 0.05. ** p<0.005

*In vitro* stimulation of our H232 and R232 STING-transfected cells lines with live vaccinia virus resulted in global downregulation of gene expression, preventing us from examining differential effects between the two STING variants. In order to avoid such confounding issues with viral infection, we stimulated cells with 2’3’ cGAMP or inactivated vaccinia virus for all further experiments.

### Gene Expression of rs1131769 Variants

mRNA was extracted from PBMCs of individuals homozygous for the CC genotype (R232) and the TT genotype (H232), and the two *TMEM173* variants were PCR amplified and cloned into lentivirus expression vectors. BJAB cells were transduced with these vectors, creating stable cell lines that constitutively overexpress each STING variant. The PCR products and completed expression vectors were both sequenced to verify the insertion of the correct genetic variants. As illustrated in Figure 4A, stable transfectants express >1,000-fold higher (and comparable between the two variants) STING mRNA than normal BJAB cells. Expression of both *TMEM173* variants transiently and minimally (less than 2-fold) increased after 2’3’ cGAMP stimulation, indicating the STING protein levels of either variant are unlikely to be significantly affected by cGAMP treatment (Figure 4B). Finally, we found that *MB21D1* (encoding cGAS, an essential upstream nucleotidyltransferase in the STING pathway that generates cyclic cGAMP) gene expression was not significantly different between the two variants, indicating that signaling pathway function upstream of STING was not affected by the STING gene variants (Figure 4C). Note that the HEK293T lines used in this report were also transfected with *MB21D1* as this cell line is known to be deficient in cGAS expression (Sun et al., 2013).

### Effect of rs1131769 Variants on Downstream Phosphorylation

Transiently transfected HEK293T and stably transfected BJAB cells were stimulated with 2’3’ cGAMP for the indicated time periods, and phosphorylation of downstream intermediates TBK1 and IRF3 was evaluated (Figure 5A & 5B). In transiently transfected HEK293T cells (also expressing cGAS), H232 STING expression was accompanied by a delay in phosphorylation of both TBK1 and IRF3 until 1 hour after stimulation. In the stably transduced BJAB line, the delayed phosphorylation was observed with both STING variants, however the magnitude of TBK1 and IRF3 phosphorylation was significantly reduced in cells expressing H232 compared to cells expressing R232 STING. No major differences in STING protein levels, or in the unphosphorylated forms of either TBK or IRF3, were observed between the cells expressing the two STING variants at the observed timepoints.

**Figure 5.**
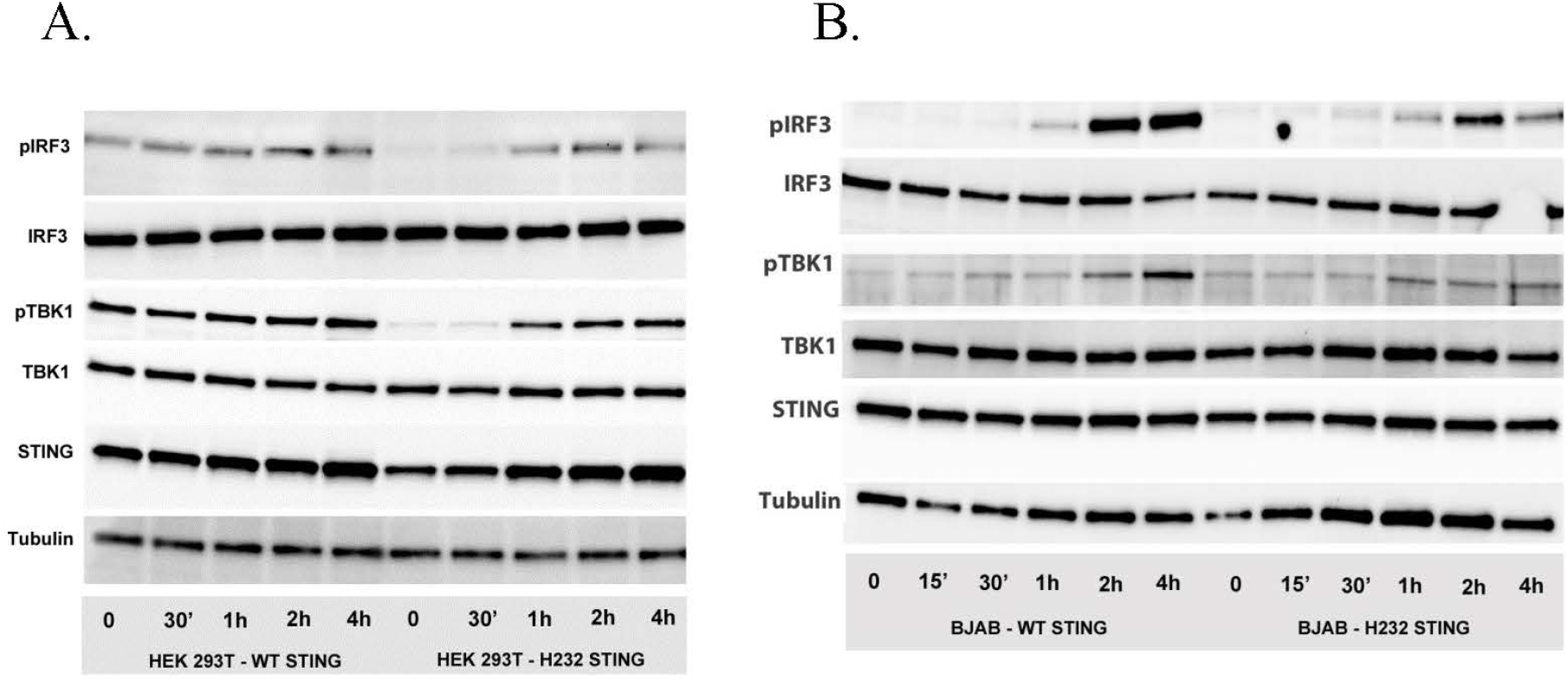
Phosphorylation of IRF3 and TBKl is delayed and decreased in the presence of H232 STING. **A)** STING pathway activation (phosphorylation of IRF3 and TBKl) after 2’3’ cGAMP stimulation of HEK 293 T cells, transiently expressing WT or H232 STING variants/alleles and cGAS. **B)** STING pathway activation (phosphorylation of IRF3 and TBKl) after 2’3’ cGAMP stimulation of lentivirus-created BJAB cell lines, stably expressing WT or H232 STING variants under Blasticidin selection.

### Effect of rs1131769 Variants on IFN Response

Reasoning that differences in TBK1 and IRF3 phosphorylation between these variants should have downstream consequences, we decided to examine differential pathway activity mediated by the two STING variants. We stimulated R232 and H232 STING variant-expressing stable BJAB cell lines with 2’3’ cGAMP and measured gene expression (qPCR, Figure 6A) and protein secretion (ELISA, Figure 6B) over time. cGAMP stimulation induced both type I (IFNα, IFNβ) and type III (IFNλ1) interferons, with significantly higher levels of IFNs (mRNA and protein) observed in R232 cells. This effect was consistent regardless of the cGAMP isomer used for stimulation (Supplemental Figure S1). Furthermore, the expression of the classical antiviral ISGs, *MX1* and *OAS1,* after 2’3’cGAMP stimulation confirmed the greater STING pathway activation in R232 STING-expressing cells over the H232 STING-expressing cells (Figure 6C).

**Figure 6.**
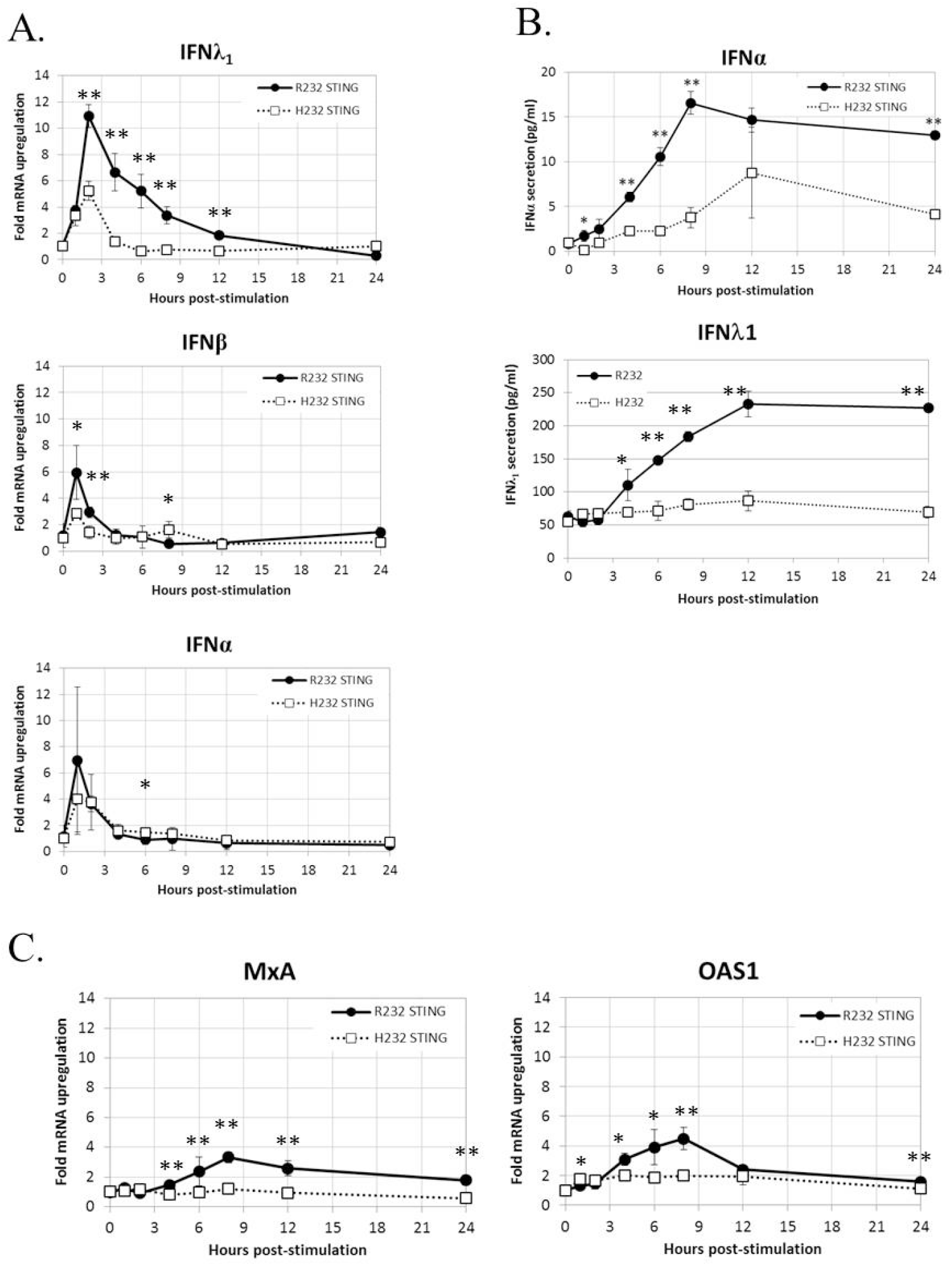
WT and H232 STING-mediated IFN response in stably transduced BJAB cells. **A)** Time course of *IFNA, IFNB, IFNL1* gene expression after cGAMP stimulation. **B)** Time course of cytokine secretion after cGAMP stimulation. **C)** Activation of representative interferon-stimulated genes after cGAMP stimulation. All data points are means and error bars representing the standard deviations of eight replicates. * p< 0.05. ** p<0.005

### Molecular Modeling of STING Variants

In order to begin elucidating the mechanism underlying the greater STING activity in R232-expressing cells, we used molecular modeling and molecular dynamics simulations to examine structural and functional differences between the R232 and H232 variants. H232 exhibited larger overall deviation from the initial experimental structure, as quantified by RMSD (Figure 7A). This greater deviation (a reflection of mobility) occurred both in the presence or absence of ligand. As RMSD is a global measure, we also quantified per-residue mobility using RMSF (Figure 7B), which indicated that the ligand-binding loops were more mobile for H232 compared to R232, particularly when the 2’3’ cGAMP ligand was bound. We further quantified the displacement of the ligand-binding loops using simple distance measures between residue 232 in each monomer. Regardless of the presence or absence of cGAMP, the ligand-binding loops of H232 were further separated from each other (Figure 7C) and from the base of the ligand-binding site (Figure 8) compared to R232. Our initial simulations assumed that the ligand-binding loops were closed over the base of the ligand-binding site; structures displayed in Figures 7D and 7E highlight the difference in STING conformation between H232 and R232. Simulations assuming an open ligand-binding loop conformation observed the same effect of H232 compared to R232 (Supplemental Figure S2). In summary, H232 exhibited greater structural flexibility and mobility of the ligand-binding loops in both the open or closed conformations and in the presence or absence of cGAMP.

**Figure 7.**
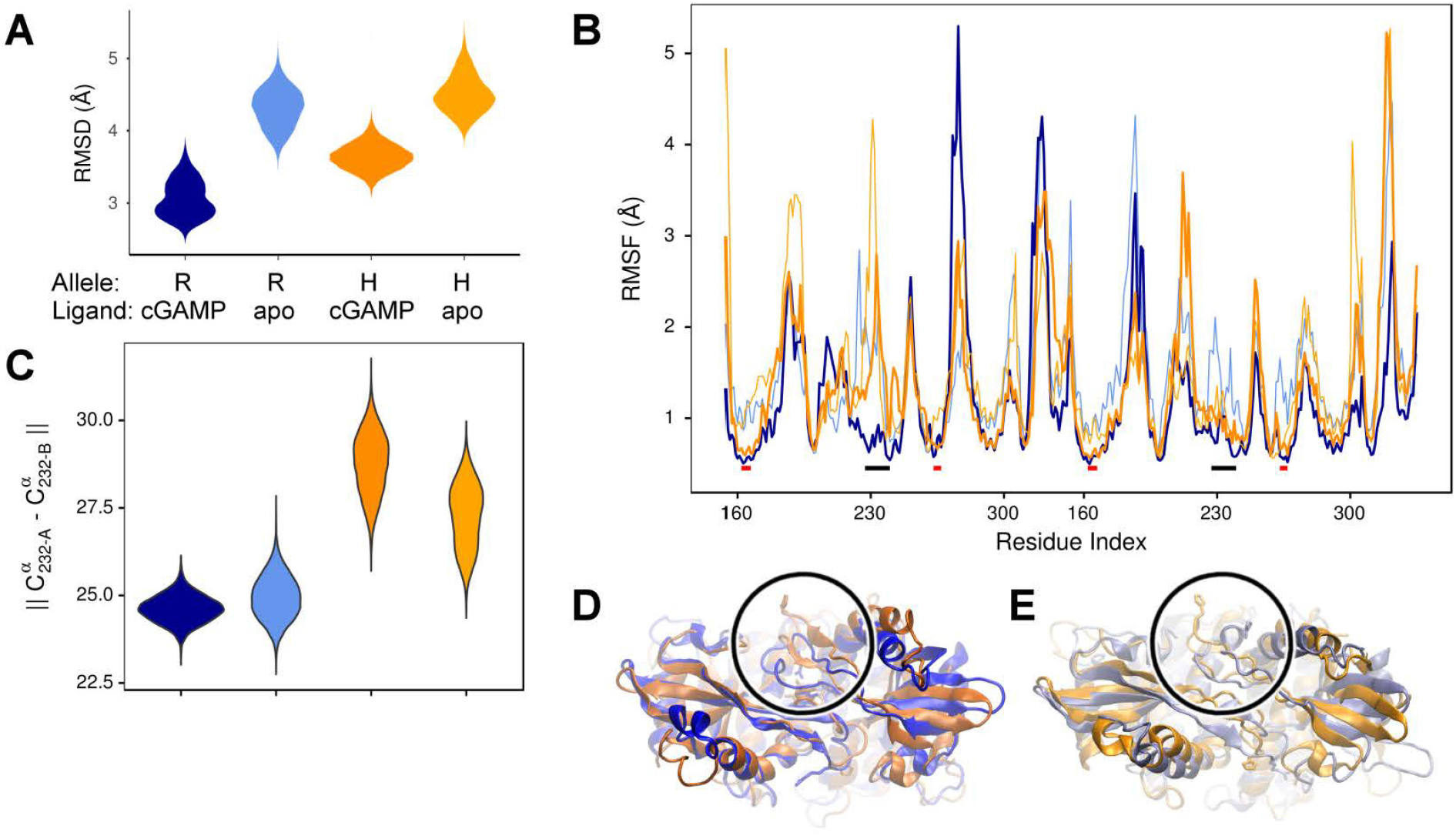
Molecular Simulation ligand binding loop contact with cGAMP in R232 (wt) and H232 STING. **A)** H232 consistently showed greater deviations from the initial open conformation, both in the presence and absence (marked as: ‘ apo’) of cGAMP. Simulation data from R232 is shown in blue, H232 in orange, and cGAMP-bound forms are in a darker shade. This color coding is continued throughout each panel. **B)** Residues within the ligand binding loops, indicated by black bars, show less difference in mobility between unbound and cGAMP-bound forms. The ligand binding site residues, indicated by red bars, are more comparable in their mobility. **C)** We monitored the distance between residue 232 from each monomer of the STING dimer as a measure of the separation of the ligand binding loops starting from the closed loop conformation. The separation was greater for H232, compared to R232, in both the unbound and cGAMP-bound forms. **D)** Representative conformations from the end of cGAMP-bound simulations are shown and the ligand binding loop circled and residue 232 (shown in ball-and-stick representation). The altered conformation of H232 is evident. **E)** Similar changes to the ligand binding loop conformation were observed in the absence of cGAMP.

**Figure 8.**
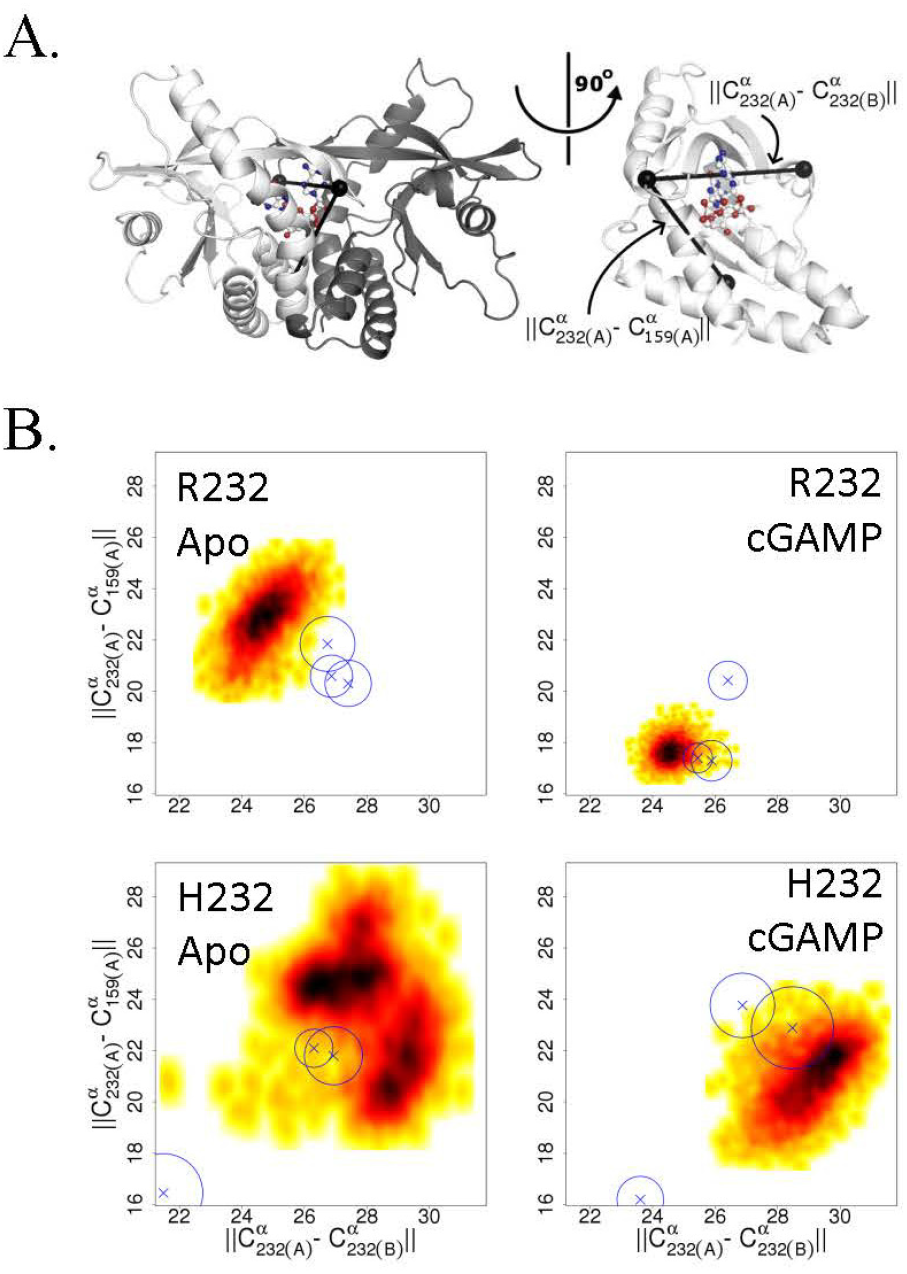
Changes in the ligand binding loop between R232 and H232 are consistent across conditions. **A)** Diagram of the STING structure showing one monomer of the closed conformation in white and the second monomer in gray, with the three C^α^ atoms used to compute distances marked by black spheres. The bound cGAMP is shown as a ball-and-stick representation. A rotated view, with the second monomer omitted for clarity, more clearly shows the relationship between these C^α^ positions and the ligand binding site. **B)** H232 lead to a consistent expansion of the distances between the ligand binding loops and from the ligand binding loops to the base of the ligand binding site. Probability density plots are shown with the most frequently sampled regions colored black and scaling through red and yellow as data become more sparse. The medians for additional independent replicates are indicated by Xs and a circle indicating the level of variability.

## Discussion

Our GWAS across two cohorts of smallpox vaccine recipients, totaling just over 1,600 individuals, identified a highly significant (p < 1 × 10^−30^) association signal from a region on chromosome 5 that was linked to significant inter-individual variations in IFNα response to *in vitro* stimulation with vaccinia virus. Fine-mapping analysis identified a number of putative causal SNPs, including several in *TMEM173*, which encodes for the signaling adaptor protein STING. STING mediates IFN responses to dsDNA and cyclic dinucleotides through a pathway involving cGAS and the phosphorylation of TBK1 and IRF3. Our regression modeling also indicated that multiple SNPs within *TMEM173* have independent effects on the phenotypic outcome. We did not identify any SNPs in other STING pathway genes associated with variations in IFNα secretion. Homozygotes for the H232 allele of rs1131769 in *TMEM173* exhibit a 90% reduction in IFNα secretion compared to R232 homozygotes. This is a highly significant and large effect that may have significant downstream consequences for poxvirus immunity. We have previously reported that a small percentage of smallpox vaccine recipients have impaired innate immune responses to vaccinia virus and that these same individuals also have suboptimal cellular immunity (Kennedy et al., 2016). Our current results provide additional support to existing data (Quigley et al., 2009; Zhao et al., 2009) suggesting that appropriate innate immune responses are necessary for robust adaptive immunity to vaccinia virus. Further investigation of this effect on smallpox immunity is warranted.

We conducted a series of experiments with the intention of elucidating functional effects of this SNP that might be underlying the identified genotype-phenotype association. We assessed gene expression of both variants of *TMEM137* by PCR and STING expression by western blot and did not detect significant differences between variants in either gene or protein expression. We observed higher promoter activity of downstream IFN-inducible genes in the R232 variant compared to the H232 variant, which suggests that there are differences in activation of the STING pathway. Upon stimulation with 2’3’ cGAMP, the R232 STING variant elicited faster phosphorylation of both TBK1 and IRF3 as well as resulted in a greater quantity of phosphorylated TBK1 and IRF3 in the cells. These changes led to a significant increase in IFN and IFN-stimulated gene expression in R232-expressing cells, confirming that the statistical association was rooted in differential biological activity. As is true of most transfection systems, the *TMEM173* gene was overexpressed in our cell lines, with the HEK293T cells expressing ~10,000 times as much *TMEM173* as untransfected cells. The BJAB transfectants also expressed high levels of *TMEM173,* but the overexpression was an order of magnitude lower. More relevant to our results, protein expression was similar to the expression levels of endogenous IRF3, TBK1, and tubulin, suggesting that protein expression was within normal limits despite the upstream overexpression of STING observed at the gene level. We note that the expression (at the gene and protein level) of both variants was consistent; therefore, differences in activity are not a result of differential gene or protein expression between the variants.

Multiple genetic variants of *TMEM173* have been described, including three non-synonymous SNPs: rs1131769 (H232R), which was the focus of our study; rs11554776 (R71H); and rs7380824 (R293Q). R71H and R293Q, together with a fourth SNP, rs78233829 (G230A), form the *HAQ* haplotype (Yi et al., 2013). Zhang et al. have previously reported that expression of the H232 variant results in reduced IFNβ transcription (Zhang et al., 2013). Our results confirm and extend these initial findings, demonstrating that IFNα secretion is also affected, as is the expression of multiple interferon-stimulated genes. Regarding the 3 alleles in the HAQ haplotype, there has been some controversy over the biological effects of these alleles (Patel et al., 2017b; a). In one study, cells carrying the G230 variant had fully functional STING activity and the HAQ haplotype effect was attributed to the R71H and R293Q SNPs (Jin et al., 2011). A similar study evaluating *TMEM173* variants found that the R232H, R293Q, and *AQ* (G230A, R293Q) variants had minimal effects on endogenous STING activity while the reduced STING function of the *HAQ* (R71H, G230A, R293Q) haplotype was attributed to the R71H variant (Yi et al., 2013). Our analysis supported previous findings that possession of the HAQ haplotype leads to reduced STING activity, conclusively demonstrated that the H232 variant also leads to reduced STING activity, and verified that this functional effect is independent of the *HAQ* haplotype. Thus, our haplotype and regression model results indicate that multiple SNPs/haplotypes are independently associated with variations in IFNα secretion. It is important to note that our study also provides a potential biochemical mechanism for the reduced IFNα activity mediated by the R232H variant. Further work will be required to tease apart the contributions of each individual SNP to the resulting immune response phenotype.

The crystal structure of STING has been resolved, as has the structure of the H232 variant bound to cGAMP (Gao et al., 2013). Our molecular modeling simulations, using these structure data, revealed that the ligand-binding loops of STING were more mobile for H232 compared to WT; that is, H232 loop conformations were more open and flexible than R232, even in the presence of bound cGAMP. We speculate that this may reflect a failure of the ligand-binding loops to either stay closed when beginning from a closed conformation, or to close when beginning from an open conformation. This may be indicative of weaker binding (and/or faster disassociation) between cGAMP and the H232 variant of STING. Our data suggest that the alterations in loop dynamics and weaker affinity of H232 STING for its ligand are the underlying molecular mechanism for the reduced STING activity that we observed for this variant.

With regard to poxviruses, it has been demonstrated that Modified Vaccinia Ankara (MVA) infection of dendritic cells (DCs) triggers type I IFN production through a STING and cGAS-dependent pathway involving cyclic dinucleotides (Dai et al., 2014). This type I IFN response is not seen during wild type vaccinia virus infection, likely due to the presence of viral immunomodulatory proteins such as C6L, E3L, and N1L in the wild type virus but not in MVA (Unterholzner et al., 2011; Cao et al., 2012). A recent report confirmed that MVA activated IRF3 in a cGAS- and STING-dependent manner, whereas wild type vaccinia strains failed to do so (Georgana et al., 2018). Our experiments found clear differences in R232 and H232 STING activity in the presence of cyclic dinucleotides and inactivated vaccinia virus. It is possible that possession of the H232 STING variant may alter the effects of viral immunomodulation of this innate immune pathway during infection. This may happen through differential interactions with viral proteins, or indirectly as reduced secretion of type I IFNs may render viral immunomodulation more effective.

We have demonstrated that carriage of the H variant of rs1131769 results in a 90% decrease in innate immune response (secreted IFNα) to vaccinia virus. This data helps resolve prior conflicting reports regarding functional effects of STING polymorphisms. We hypothesize that the effect of this polymorphism is due to different flexibility/mobility in STING H232 loop conformations, which results in reduced ability of H232 STING to phosphorylate downstream signaling intermediates and mediate effective STING pathway activation.

Poxviruses represent a continuing public health concern due to the risk of bioterrorism use, zoonotic outbreaks (e.g., monkeypox, buffalopox, vaccinia-like viruses, and novel poxviruses), the increasing use of poxviruses for oncolytic viral therapy, and their use as vectors for vaccine antigens against HIV, rabies, Ebola, Zika, and other pathogens. STING also plays an essential role in triggering protective innate responses to DNA viruses (e.g., poxviruses, herpes simplex viruses, varicella zoster, EBV, HPV, and others) and multiple bacterial pathogens. Polymorphisms that reduce the effectiveness of the innate response to these threats are likely to enhance disease susceptibility and may have a deleterious effect on vaccine immunogenicity in the ~15% of the population with this genotype. Given the broad potential impact of this pathway, this is an area that merits additional investigation.

Understanding how genetic factors control the immune response to poxviruses will have important clinical implications in how, when, and in whom these vectors can be safely and effectively used. Furthermore, this information may inform the use of adjuvants to overcome this defect and enhance vaccine responses or the development of therapeutic drugs that can be used to enhance the innate antiviral response during an infection.

## Author contributions

**RBK** contributed to the conception and design of the study, participated in the acquisition of data as well as the analysis and interpretation of the results. He prepared the initial draft of the manuscript and revised it for intellectual content.

**IHH** contributed to the study design, participated in data acquisition and interpretation of study results. She assisted in drafting the manuscript, revised it for intellectual content and approved the final version.

**IGO** contributed to the study design, participated in data acquisition and interpretation of study results. She assisted in drafting the manuscript, revised it for intellectual content and approved the final version.

**EAV** contributed to the study design, participated in data acquisition and interpretation of study results. She assisted in drafting the manuscript, revised it for intellectual content and approved the final version.

**BRL** contributed data analysis and interpretation. She assisted in drafting the manuscript, revised it for intellectual content and approved the final version.

**DJS** supervised and contributed to the data analysis and interpretation. He participated in drafting and revising the manuscript, and approved the final version.

**MTZ** contributed to the interpretation of study results, molecular modeling, data analysis, drafting the manuscript, critical review, and approved the final version.

**ALO** contributed to the design of the study, participated in data analysis and interpretation, assisted in drafting the manuscript, revised the manuscript for intellectual content, and approved the final version.

**GAP** contributed to the conception and design of the study, participated in data analysis and interpretation, secured funding for the project, participated in drafting and revising the manuscript, and approved the final version.

## Acknowledgements

We gratefully thank the study subjects for their participation. We also thank Drs. Megan Ryan and Kevin Russell, as well as the clinical staff at the Naval Health Research Center for assistance with subject recruitment. We thank Nathaniel D. Warner and Krista M. Goergen for statistical and programming assistance and Caroline L. Vitse for proofreading and editorial assistance. This study was supported by NIH through the NIAID Population Genetics Analysis Program Contract No.HHSN266200400065C and Contract No. HHSN272201000025C, and by the National Center for Research Resources grant 1 UL1 RR024150-01. The content is solely the responsibility of the authors and does not necessarily represent the official views of the National Institutes of Health.

The study data are all available, without restriction, at ImmPort. https://immport.niaid.nih.gov Study #:SDY28

## Competing interests

Dr. Poland is the chair of a Safety Evaluation Committee for novel investigational vaccine trials being conducted by Merck Research Laboratories. Dr. Poland offers consultative advice on vaccine development to Merck & Co. Inc., Avianax, Adjuvance, Valneva, Medicago, Sanofi Pasteur, GlaxoSmithKline, Emergent Biosolutions, and Dynavax. Drs. Kennedy, Poland, and Ovsyannikova hold three patents related to vaccinia and measles peptide research. Dr. Kennedy has received funding from Merck Research Laboratories to study waning immunity to mumps vaccine. These activities have been reviewed by the Mayo Clinic Conflict of Interest Review Board and are conducted in compliance with Mayo Clinic Conflict of Interest policies. This research has been reviewed by the Mayo Clinic Conflict of Interest Review Board and was conducted in compliance with Mayo Clinic Conflict of Interest policies.

## Notes

https://immport.niaid.nih.gov

