## Supplemental Table 1 for "Polymorphisms in STING affect human innate immune responses to poxviruses"

**Supplemental Table 1. Smallpox GWAS Cohort Demographics**

| **Supplemental Table 1. Cohort Demographics** | |
| --- | --- |
| *Total subjects* | 1,653 |
| *Gender* |  |
| Male | 1,365 (82.6%) |
| Female | 288 (17.4%) |
| *Age* | 19 – 41 years old |
| *Ethnicity* |  |
| Hispanic/Latino | 381 (23%) |
| Not Hispanic/Latino | 1248 (75.5%) |
| Unknown | 24 (1.5%) |
| *Race* |  |
| American Indian/Alaska Native | 26 (1.6%) |
| Asian, Native Hawaiian, Pacific Islander | 12 (0.7%) |
| African American | 25 (1.5%) |
| Caucasian | 1,351 (81.7%) |
| More than one race | 81 (4.9%) |
| Other | 120 (7.3%) |
| Unknown | 38 (2.3%) |
| *Time since immunization* |  |
| Median | 1.7 years |
| Range | 1.1 – 2.8 years |
