## Supplementary figures and images for "Polymorphisms in STING affect human innate immune responses to poxviruses"

### Supplemental Figure 1

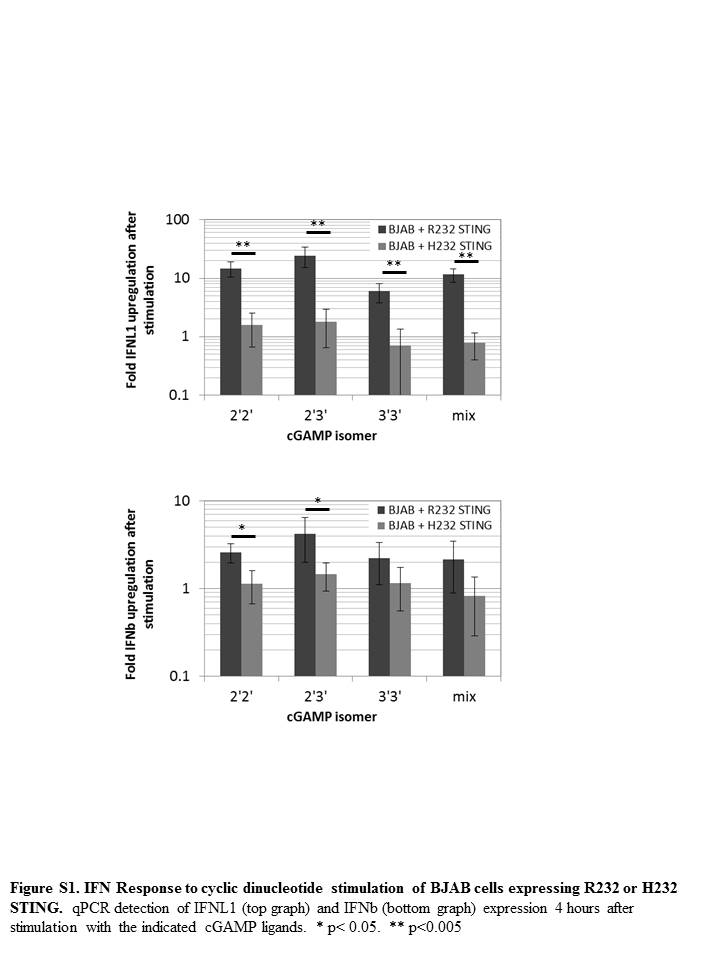

### Supplemental Figure 2

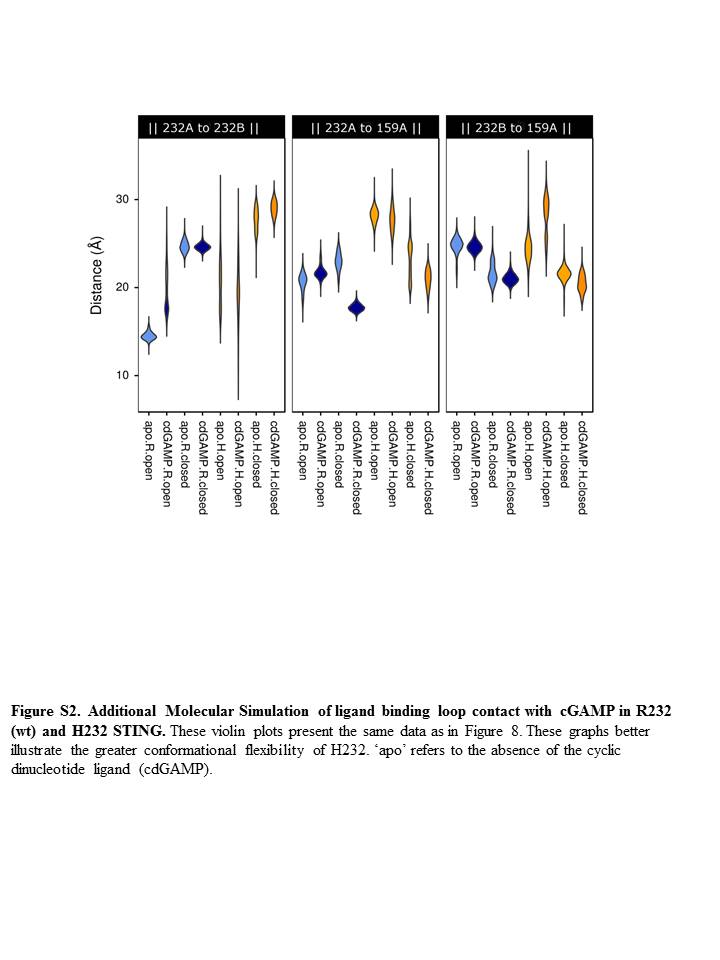
